# GraphDTA: Predicting drug–target binding affinity with graph neural networks

**DOI:** 10.1101/684662

**Authors:** Thin Nguyen, Hang Le, Thomas P. Quinn, Tri Nguyen, Thuc Duy Le, Svetha Venkatesh

## Abstract

The development of new drugs is costly, time consuming, and often accompanied with safety issues. Drug repurposing can avoid the expensive and lengthy process of drug development by finding new uses for already approved drugs. In order to repurpose drugs effectively, it is useful to know which proteins are targeted by which drugs. Computational models that estimate the interaction strength of new drug--target pairs have the potential to expedite drug repurposing. Several models have been proposed for this task. However, these models represent the drugs as strings, which is not a natural way to represent molecules. We propose a new model called **GraphDTA** that represents drugs as graphs and uses graph neural networks to predict drug--target affinity. We show that graph neural networks not only predict drug--target affinity better than non-deep learning models, but also outperform competing deep learning methods. Our results confirm that deep learning models are appropriate for drug--target binding affinity prediction, and that representing drugs as graphs can lead to further improvements.

**Availability of data and materials:** The proposed models are implemented in Python. Related data, pre-trained models, and source code are publicly available at https://github.com/thinng/GraphDTA. All scripts and data needed to reproduce the post-hoc statistical analysis are available from https://doi.org/10.5281/zenodo.3603523.

## 1 Background

It costs about 2.6 billion US dollars to develop a new drug [1], and can take up to 17 years for FDA approval [2, 3]. Finding new uses for already approved drugs avoids the expensive and lengthy process of drug development [2, 4]. For example, nearly 70 existing FDA-approved drugs are currently being investigated to see if they can be repurposed to treat COVID-19 [5]. In order to repurpose drugs effectively, it is useful to know which proteins are targeted by which drugs. High-throughput screening experiments are used to examine the affinity of a drug toward its targets; however, these experiments are costly and time-consuming [6, 7], and an exhaustive search is infeasible because there are millions of drug-like compounds [8] and hundreds of potential targets [9, 10]. As such, there is a strong motivation to build computational models that can estimate the interaction strength of new drug–target pairs based on previous drug–target experiments.

Several computational approaches have been proposed for drug–target affinity (**DTA**) prediction [11, 12, 13]. One approach is molecular docking, which predicts the stable 3D structure of a drug-target complex via a scoring function [14]. Even though the molecular docking approach is potentially more informative, it require knowledge about the crystallized structure of proteins which may not be available. Another approach uses collaborative filtering. For example, the **SimBoost** model uses the affinity similarities among drugs and among targets to build new features. These features are then used as input in a gradient boosting machine to predict the binding affinity for unknown drug–target pairs [15]. Alternatively, the similarities could come from others sources (rather than the training data affinities). For example, kernel-based methods use kernels built from molecular descriptors of the drugs and targets within a regularized least squares regression (RLS) framework [16, 17]. To speed up model training, the **KronRLS** model computes a pairwise kernel *K* from the Kronecker product of the drug-by-drug and protein-by-protein kernels [16, 17] (for which any similarity measure can be used). **DTA** prediction may also benefit from adopting methods for predicting drug–target interactions (**DTI**). Approaches in this line of work include DTI-CDF [18], a cascade deep forest model, or DTI-MLCD [19], a multi-label learning supported with community detection.

Another approach uses neural networks trained on 1D representations of the drug and protein sequences. For example, the **DeepDTA** model uses 1D representations and layers of 1D convolutions (with pooling) to capture predictive patterns within the data [20]. The final convolution layers are then concatenated, passed through a number of hidden layers, and regressed with the drug–target affinity scores. The **WideDTA** model is an extension of **DeepDTA** in which the sequences of the drugs and proteins are first summarized as higher-order features [21]. For example, the drugs are represented by the most common sub-structures (the Ligand Maximum Common Substructures (LMCS) [22]), while the proteins are represented by the most conserved sub-sequences (the Protein Domain profiles or Motifs (PDM) from PROSITE [23]). While **WideDTA** [21] and **DeepDTA** [20] learn a latent feature vector for each protein, the **PADME** model [24] uses fixed-rule descriptors to represent proteins, and performs similarly to **DeepDTA** [20].

The deep learning models are among the best performers in DTA prediction [25]. However, these models represent the drugs as strings, which are not a natural way to represent molecules. When using strings, the structural information of the molecule is lost, which could impair the predictive power of a model as well as the functional relevance of the learned latent space. Already, graph convolutional networks have been used in computational drug discovery, including interaction prediction, synthesis prediction, *de novo* molecular design, and quantitative structure prediction [26, 27, 28, 29, 30, 31, 25]. However, graph neural networks have not been used for DTA prediction. Of these, [26, 25, 32] are closest to our work, but look at binary prediction, while our model looks to predict a continuous value of binding affinity. Also, in [25], the input is a drug descriptor (single input), while our model takes as input both a drug descriptor and a sequence (dual input).

In this article, we propose **GraphDTA**, a new neural network architecture capable of directly modelling drugs as molecular graphs, and show that this approach outperforms state-of-the-art deep learning models on two drug–target affinity prediction benchmarks. The approach is based on the solution we submitted to the IDG-DREAM Drug-Kinase Binding Prediction Challenge^1^, where we were among the Top Ten Performers from 530 registered participants^2^. In order to better understand how our graph-based model works, we performed a multivariable statistical analysis of the model’s latent space. We identified correlations between hidden node activations and domain-specific drug annotations, such as the number of aliphatic OH groups, which suggests that our graph neural network can automatically assign importance to well-defined chemical features without any prior knowledge. We also examine the model’s performance and find that a handful of drugs contribute disproportionately to the total prediction error, and that these drugs are inliers (i.e., not outliers) in an ordination of the model’s latent space. Taken together, our results suggest that graph neural networks are highly accurate, abstract meaningful concepts, and yet fail in predictable ways. We conclude with a discussion about how these insights can feedback into the research cycle.

## 2 Methods

### 2.1 Overview of GraphDTA

We propose a novel deep learning model called **GraphDTA** for drug–target affinity (**DTA**) prediction. We frame the DTA prediction problem as a regression task where the input is a drug–target pair and the output is a continuous measurement of binding affinity for that pair. Existing methods represent the input drugs and proteins as 1D sequences. Our approach is different; we represent the drugs as molecular graphs so that the model can directly capture the bonds among atoms.

### 2.2 Drug representation

**SMILES** (Simplified Molecular Input Line Entry System) was invented to represent molecules to be readable by computers [33], enabling several efficient applications, including fast retrieval and substructure searching. From the SMILES code, drug descriptors like the number of heavy atoms or valence electrons can be inferred and readily used as features for affinity prediction. One could also view the SMILES code as a string. Then, one could featurize the strings with natural language processing (NLP) techniques, or use them directly in a convolutional neural network (CNN).

Instead, we view drug compounds as a graph of the interactions between atoms, and build our model around this conceptualization. To describe a node in the graph, we use a set of atomic features adapted from **DeepChem** [34]. Here, each node is a multi-dimensional binary feature vector expressing five pieces of information: the atom symbol, the number of adjacent atoms, the number of adjacent hydrogens, the implicit value of the atom, and whether the atom is in an aromatic structure [34]. We convert the SMILES code to its corresponding molecular graph and extract atomic features using the open-source chemical informatics software **RDKit** [35].

### 2.3 Protein representation

One-hot encoding has been used in previous works to represent both drugs and proteins, as well as other biological sequences like DNA and RNA. This paper tests the hypothesis that a graph structure could yield a better representation for drugs, and so only drugs were represented as a graph. Although one could also represent proteins as graphs, doing so is more difficult because the tertiary structure is not always available in a reliable form. As such, we elected to use the popular one-hot encoding representation of proteins instead.

For each target in the experimented datasets, a protein sequence is obtained from the UniProt database using the target’s gene name. The sequence is a string of ASCII characters which represent amino acids. Each amino acid type is encoded with an integer based on its associated alphabetical symbol (e.g., Alanine (A) is 1, Cystine (C) is 3, Aspartic Acid (D) is 4, and so on), allowing the protein to be represented as an integer sequence. To make it convenient for training, the sequence is cut or padded to a fixed length sequence of 1000 residues. In case a sequence is shorter, it is padded with zero values.

These integer sequences are used as input to the embedding layers which return a 128-dimensional vector representation. Next, three 1D convolutional layers are used to learn different levels of abstract features from the input. Finally, a max pooling layer is applied to get a representation vector of the input protein sequence.

### 2.4 Deep learning on molecular graphs

Having the drug compounds represented as graphs, the task now is to design an algorithm that learns effectively from graphical data. The recent success of CNN in computer vision, speech recognition, and natural language processing has encouraged research into graph convolution. A number of works have been proposed to handle two main challenges in generalizing CNN to graphs: (1) the formation of receptive fields in graphs whose data points are not arranged as Euclidean grids, and (2) the pooling operation to down-sample a graph. These new models are called graph neural networks.

In this work, we propose a new DTA prediction model based on a combination of graph neural networks and conventional CNN. Figure 1 shows a schematic of the model. For the proteins, we use a string of ASCII characters and apply several 1D CNN layers over the text to learn a sequence representation vector. Specifically, the protein sequence is first categorically encoded, then an embedding layer is added to the sequence where each (encoded) character is represented by a 128-dimensional vector. Next, three 1D convolutional layers are used to learn different levels of abstract features from the input. Finally, a max pooling layer is applied to get a representation vector of the input protein sequence. This approach is similar to the existing baseline models. For the drugs, we use the molecular graphs and trial 4 graph neural network variants, including GCN [36], GAT [37], GIN [38], and a combined GAT-GCN architecture, all of which we describe below.

**Figure 1:**
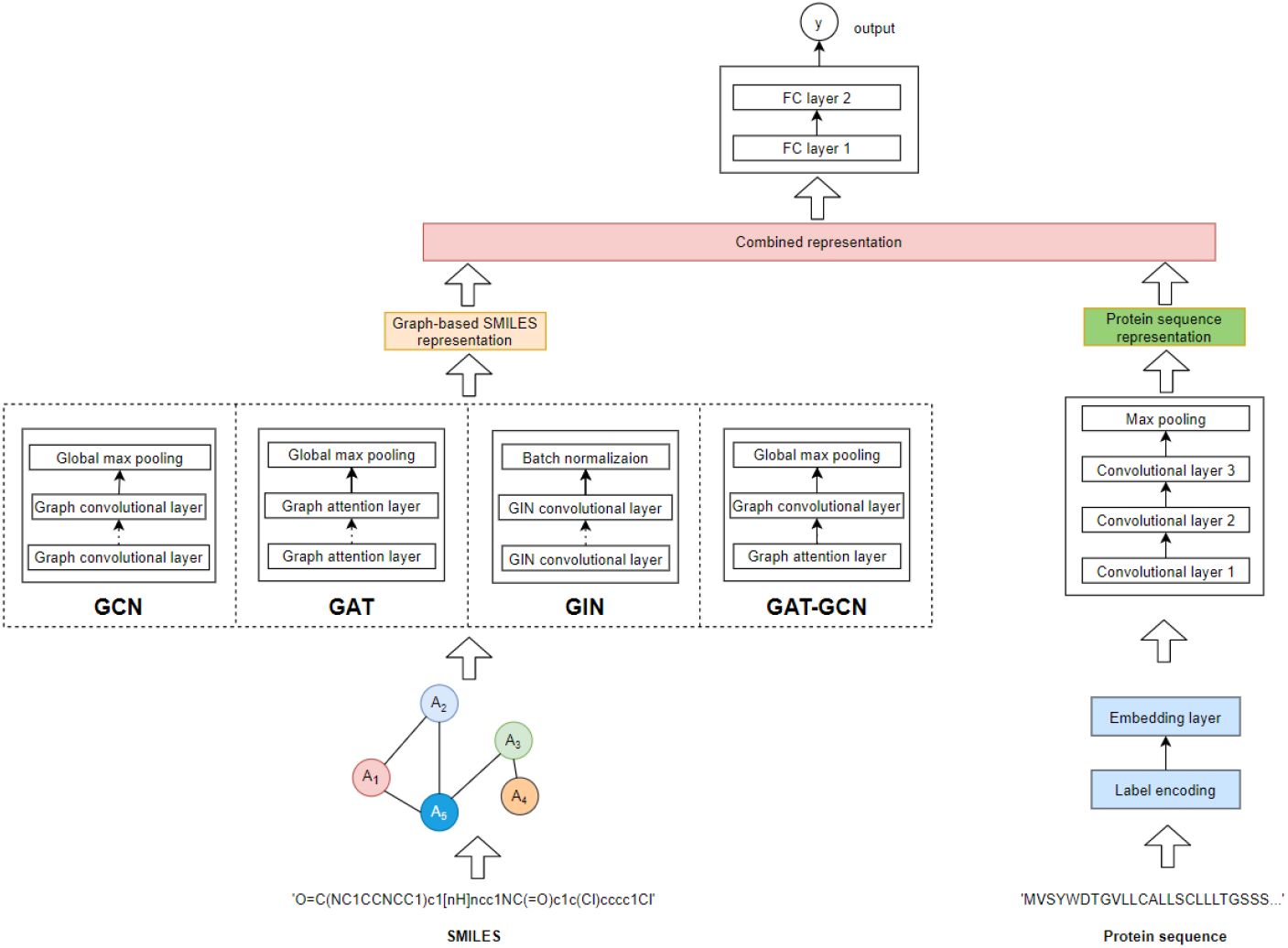
This figure shows the **GraphDTA** architecture. It takes a drug–target pair as the input data, and the pair’s affinity as the output data. It works in 3 stages. First, the SMILES code of a drug is converted into a molecular graph, and a deep learning algorithm learns a graph representation. Meanwhile, the protein sequence is encoded and embedded, and several 1D convolutional layers learn a sequence representation. Finally, the two representation vectors are concatenated and passed through several fully connected layers to estimate the output drug–target affinity value.

#### 2.4.1 Variant 1: GCN-based graph representation learning

In this work, we focus on **predicting a continuous value** indicating the level of interaction of a drug and a protein sequence. Each drug is encoded as a graph and each protein is represented as a string of characters. To this aim, we make use of GCN model [36] for learning on graph representation of drugs. Note that, however, the original GCN is designed for semi-supervised node classification problem, i.e., the model learns the **node-level feature vectors**. For our goal, to estimate the drug-protein interaction, a **graph-level representation** of each drug is required. Common techniques to aggregate the whole graph feature from learned node features include Sum, Average, and Max Pooling. In our experiments, the use of Max Pooling layer in GCN-based GraphDTA usually results in better performance compared to that of the remaining.

Formally, denote a graph for a given drug as *G* = (*V, E*), where *V* is the set of *N* nodes each is represented by a *C*-dimensional vector and *E* is the set of edges represented as an adjacency matrix *A*. A multi-layer graph convolutional network (GCN) takes as input a node feature matrix *X* ∈ ℝ^*N*×*C*^ (*N* = |*V*|, *C*: the number of features per node) and an adjacency matrix *A* ∈ ℝ^*N*×*N*^; then produces a node-level output *Z* ∈ ℝ^*N*×*F*^ (*F* : the number of output features per node). A propagation rule can be written in the normalized form for stability, as in [36]:

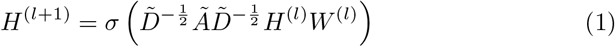

where *Ã* = *A* + *I*_*N*_ is the adjacency matrix of the undirected graph with added self-connections, 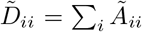; *H*^(*l*)^ ∈ ℝ^*N*×*C*^ is the matrix of activation in the *l*^*th*^ layer, *H*^(0)^ = *X*, *σ* is an activation function, and *W* is learnable parameters.

A layer-wise convolution operation can be approximated, as in [36]:

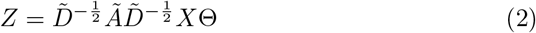

where Θ ∈ ℝ^*C*×*F*^ (*F* : the number of filters or feature maps) is the matrix of filter parameters.

Note that, however, the GCN model learns node-level outputs *Z* ∈ ℝ^*N*×*F*^. To make the GCN applicable to the task of learning a representation vector of the whole graph, we add a global max pooling layer right after the last GCN layer. In our GCN-based model, we use three consecutive GCN layers, each activated by a ReLU function. Then a global max pooling layer is added to obtain the graph representation vector.

#### 2.4.2 Variant 2: GAT-based graph representation learning

Unlike graph convolution, the graph attention network (**GAT**) [37] proposes an attention-based architecture to learn hidden representations of nodes in a graph by applying a self-attention mechanism. The building block of a GAT architecture is a *graph attention layer*. The GAT layer takes the set of nodes of a graph as input, and applies a linear transformation to every node by a weigh matrix **W**. For each input node *i* in the graph, the *attention coefficients* between *i* and its first-order neighbors are computed as

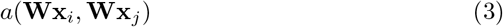

This value indicates the importance of node *j* to node *i*. These *attention coefficients* are then normalized by applying a soft-max function, then used to compute the output features for nodes as

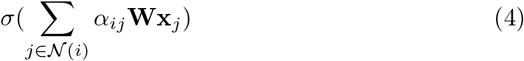

where *σ*(.) is a non-linear activation function and *α*_*ij*_ are the normalized *attention coefficients*.

In our model, the GAT-based graph learning architecture includes two GAT layers, activated by a ReLU function, then followed a global max pooling layer to obtain the graph representation vector. For the first GAT layer, *multi-head-attentions* are applied with the number of heads set to 10, and the number of output features set to the number of input features. The number of output features of the second GAT is set to 128.

#### 2.4.3 Variant 3: Graph Isomorphism Network (GIN)

The graph isomorphism network (**GIN**) [38] is newer method that supposedly achieves maximum discriminative power among graph neural networks. Specifically, GIN uses a multi-layer perceptron (MLP) model to update the node features as

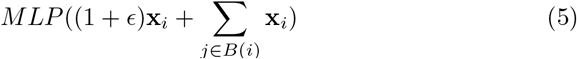

where *∊* is either a learnable parameter or a fixed scalar, **x** is the node feature vector, and *B*(*i*) is the set of nodes neighboring *i*.

In our model, the GIN-based graph neural net consists of five GIN layers, each followed by a batch normalization layer. Finally, a global max pooling layer is added to obtain the graph representation vector.

#### 2.4.4 Variant 4: GAT-GCN combined graph neural network

We also investigate a combined **GAT-GCN** model. Here, the graph neural network begins with a GAT layer that takes the graph as input, then passes a convolved feature matrix to the subsequent GCN layer. Each layer is activated by a ReLU function. The final graph representation vector is then computed by concatenating the global max pooling and global mean pooling layers from the GCN layer output.

### 2.5 Benchmark

To compare our model with the state-of-the-art **DeepDTA** [20] and **WideDTA** [21] models, we use the same datasets from the [20, 21] benchmarks:

- **Davis** contains the binding affinities for all pairs of 72 drugs and 442 targets, measured as *K*_*d*_ constants and ranging from 5.0 to 10.8 [39].
- **Kiba** contains the binding affinities for 2,116 drugs and 229 targets, measured as KIBA scores and ranging from 0.0 to 17.2 [40].

To make the comparison as fair as possible, we use the same set of training and testing examples from [20, 21], as well as the same performance metrics: Mean Square Error (**MSE**, the smaller the better) and Concordance Index (**CI**, the larger the better). For all baseline methods, we report the performance metrics as originally published in [20, 21]. The hyper-parameters used for our experiments are summarized in Table 3. The hyper-parameters were not tuned, but chosen *a priori* based on our past modelling experience.

### 2.6 Model interpretation

The activation of nodes within layers of a deep neural network are called latent variables, and can be analyzed directly to understand how a model’s performance relates to domain knowledge [41]. We obtained the 128 latent variables from the graph neural network layer, and analyzed them directly through a redundancy analysis. This multivariable statistical method allows us to measure the percent of the total variance within the latent variables that can be explained by an external data source. In our case, the external data source is a matrix of 38 molecular JoeLib features/descriptors [42] for each drug (available from ChemMine Tools [43]). We also compare the value of the principal components from these latent variables with the per-drug test set error. Here, the per-drug (or per-protein) error refers to the median of the absolute error between the predicted DTA and the ground-truth DTA for all test set pairs containing that drug (or that protein). For these analyses, we focus on the GIN model [38] (because of its superior performance) and the Kiba dataset [40] (because of its larger drug catalog).

## 3 Results and Discussion

### 3.1 Graphical models outperform the state-of-the-art

Table 1 compares the performance of 4 variant **GraphDTA** models with the existing baseline models for the Davis dataset. Here, all 4 variants had the lowest MSE. The best variant had an MSE of 0.229 which is 14.0% lower than the best baseline of 0.261. The improvement is less obvious according to the CI metric, where only 2 of the 4 variants had the highest CI. The best CI for a baseline model was 0.886. By comparison, the GAT and GIN models achieved a CI of 0.892 and 0.893, respectively.

**Table 1:** Prediction performance on the Davis dataset, sorted by MSE. Baseline results are from [20, 21]. We compare 4 graph neural network variants: GIN [38], GAT [37], GCN [36], and combined GAT-GCN [37, 36]. Italics: best for baseline models, bold: better than baselines.

| Method | Protein rep. | Compound rep. | CI | MSE |
| --- | --- | --- | --- | --- |
| Baseline models |  |  |  |  |
| DeepDTA [20] | Smith-Waterman | Pubchem-Sim | 0.790 | 0.608 |
| DeepDTA [20] | Smith-Waterman | 1D | *0.886* | 0.420 |
| DeepDTA [20] | 1D | Pubchem-Sim | 0.835 | 0.419 |
| KronRLS [16, 17] | Smith-Waterman | Pubchem-Sim | 0.871 | 0.379 |
| SimBoost [15] | Smith-Waterman | Pubchem-Sim | 0.872 | 0.282 |
| DeepDTA [20] | 1D | 1D | 0.878 | *0.261* |
| WideDTA [21] | 1D + PDM | 1D + LMCS | *0.886* | 0.262 |
| Proposed method - **GraphDTA** |  |  |  |  |
| GCN | 1D | Graph | 0.880 | **0.254** |
| GAT-GCN | 1D | Graph | 0.881 | **0.245** |
| GAT | 1D | Graph | **0.892** | **0.232** |
| GIN | 1D | Graph | **0.893** | **0.229** |

Table 2 compares the performance of the **GraphDTA** models with the existing baseline models for the Kiba dataset. Here, 3 of the 4 variants had the lowest MSE and the highest CI, including GIN, GCN, and GAT-GCN. Of note, the best MSE here is 0.139, which is 28.8% lower than the best baseline of 0.179. Of all variants tested, GIN is the only one that had the best performance for both datasets and for both performance measures. For this reason, we focus on the GIN in all post-hoc statistical analyses.

**Table 2:** Prediction performance on the Kiba dataset, sorted by MSE. Baseline results are from [20, 21]. We compare 4 graph neural network variants: GIN [38], GAT [37], GCN [36], and combined GAT-GCN [37, 36]. Italics: best for baseline models, bold: better than baselines.

| Method | Protein rep. | Compound rep. | CI | MSE |
| --- | --- | --- | --- | --- |
| Baseline models |  |  |  |  |
| DeepDTA [20] | 1D | Pubchem-Sim | 0.718 | 0.571 |
| DeepDTA [20] | Smith-Waterman | Pubchem-Sim | 0.710 | 0.502 |
| KronRLS [16, 17] | Smith-Waterman | Pubchem-Sim | 0.782 | 0.411 |
| SimBoost [15] | Smith-Waterman | Pubchem-Sim | 0.836 | 0.222 |
| DeepDTA [20] | Smith-Waterman | 1D | 0.854 | 0.204 |
| DeepDTA [20] | 1D | 1D | 0.863 | 0.194 |
| WideDTA [21] | 1D + PDM | 1D + LMCS | *0.875* | *0.179* |
| Proposed method - **GraphDTA** |  |  |  |  |
| GAT | 1D | Graph | 0.866 | 0.179 |
| GIN | 1D | Graph | **0.882** | **0.147** |
| GCN | 1D | Graph | **0.889** | **0.139** |
| GAT_GCN | 1D | Graph | **0.891** | **0.139** |

**Table 3:** Hyper-parameters for different graph neural network variants used in our experiments.

| Hyper-parameters | Setting |
| --- | --- |
| Learning rate | 0.0005 |
| Batch size | 512 |
| Optimizer | Adam |
| GCN layers | 3 |
| GIN layers | 5 |
| GAT layers | 2 |
| GAT_GCN layers | 2 |

### 3.2 Graphical models discover known drug properties

A graph neural network works by abstracting the molecular graph of each drug into a new feature vector of *latent variables*. In our model, there are 128 latent variables which together characterise the structural properties of the drug. Since the latent variables are learned during the DTA prediction task, we assume that they represent graphical features that contribute meaningfully to DTA.

Unfortunately, it is not straightforward to determine the molecular substructures to which each latent variable corresponds. However, we can regress the learned latent space with a matrix of known molecular descriptors to look for overlap. Figure 2 shows a redundancy analysis of the 128 latent variables regressed with 38 molecular descriptors [42] (available from ChemMine Tools [43]). From this, we find that 20.19% of the latent space is explained by the known descriptors, with the “Number of aliphatic OH groups” contributing most to the explained variance. Indeed, two latent variables correlate strongly with this descriptor: hidden nodes V58 and V14 both tend to have high activation when the number of aliphatic OH groups is large. This finding provides some insight into how the graphical model might “see” the drugs as a set of molecular sub-structures, though most of the latent space is orthogonal to the known molecular descriptors.

**Figure 2:**
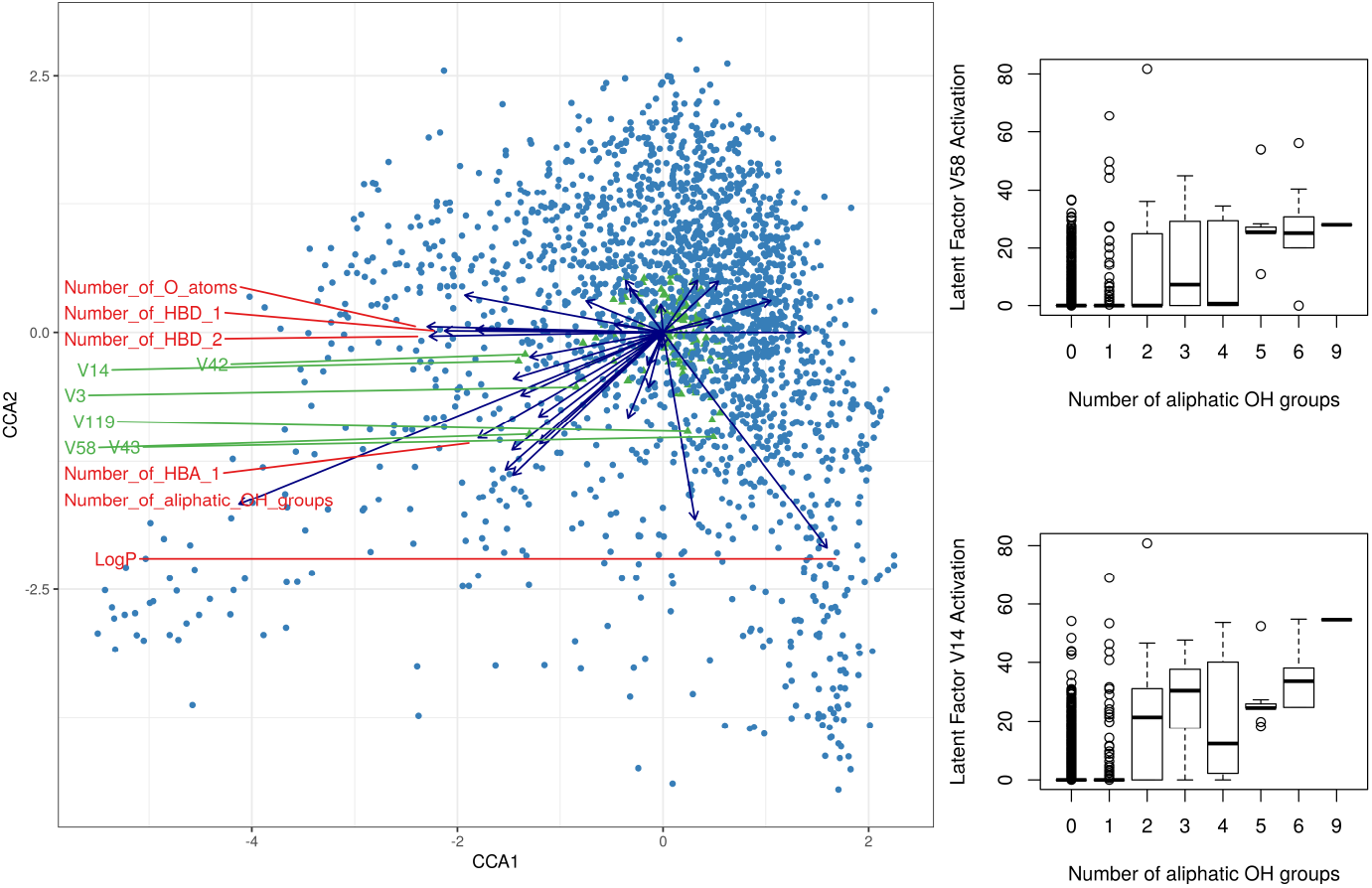
The left panel of the figure shows a redundancy analysis triplot for the 128 drug latent variables regressed with 38 JoeLib molecular descriptors [42]. The blue dots represent drugs, the green dots represent latent variables (the 6 furthest from origin are labelled), and the arrows represent molecular descriptors (the 5 longest are labelled). The right panel of the figure shows the activation of two latent variables plotted against the number of aliphatic OH groups in that drug. These results suggest that the graph convolutional network can abstract known molecular descriptors without any prior knowledge.

### 3.3 A few drugs contribute disproportionately to total error

Although the **GraphDTA** model outperforms its competitors, we wanted to know more about why its predictions sometimes failed. For this, we averaged the prediction error for each drug (and each protein), for both the Davis and Kiba test sets. Figures 3 and 4 show the median of the absolute error (MAE) for affinity prediction, sorted from smallest to largest. Interestingly, we see that a handful of drugs (and a handful of proteins) contribute disproportionately to the overall error. Of note, CHEMBL1779202 (an ALK inhibitor), CHEMBL1765740 (a PDK1 inhibitor) and the protein CSNK1E all had an MAE above 2.

**Figure 3:**
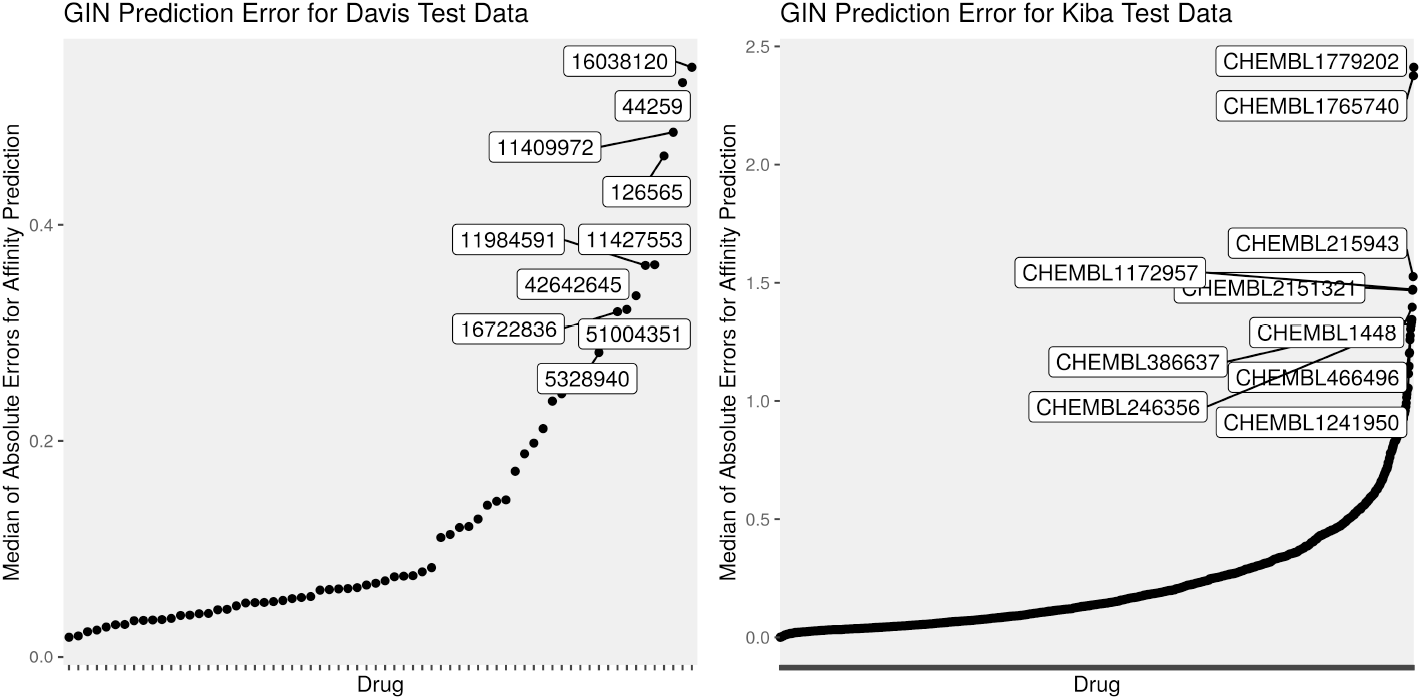
This figure shows the median of the absolute error for each drug, sorted in increasing order, for the Davis and Kiba test sets. Here, we see that the errors are not distributed evenly across the drugs. It is harder to predict the target affinities for some drugs than others.

**Figure 4:**
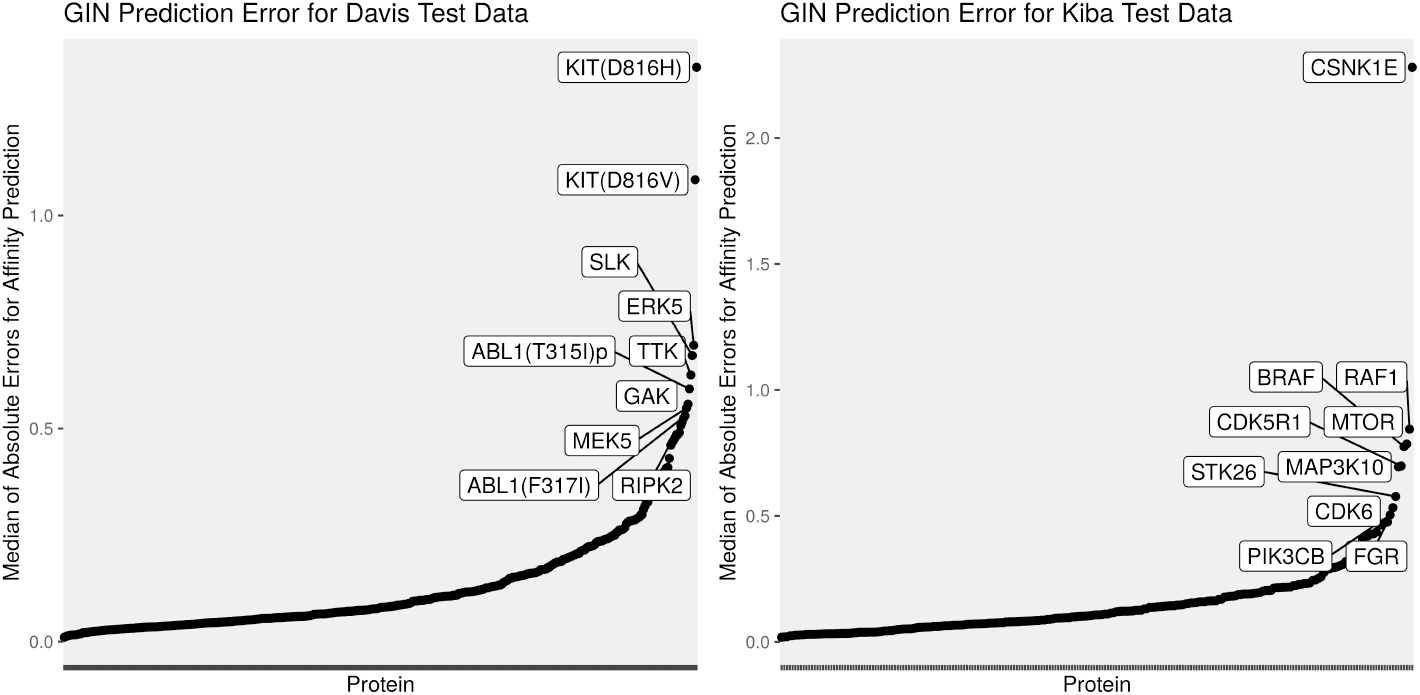
This figure shows the median of the absolute error for each protein, sorted in increasing order, for the Davis and Kiba test sets. Here, we see that the errors are not distributed evenly across the proteins. It is harder to predict the target affinities for some proteins than others.

We examined the latent space with regard to the prediction error, but could not find any obvious pattern that separated hard-to-predict drugs from easy-to-predict drugs. The only trend we could find is that the easy-to-predict drugs are more likely to appear as outliers in a PCA of the latent space. Supplemental Figure 1 (https://github.com/thinng/GraphDTA/blob/master/supplement.pdf) shows the median errors plotted against the first six principal components, where we see that the hard-to-predict drugs usually appear close to the origin. We interpret this to mean that drugs with unique molecular sub-structures are always easy to predict. On the other hand, the hard-to-predict drugs tend to lack unique structures, though this is apparently true for many easy-to-predict drugs too.

### 3.4 Model interpretation and the research cycle

Knowing how a model works and when a model fails can feedback into the research cycle. In the post-hoc statistical analysis of our model, we find that a graph neural network can learn the importance of known molecular descriptors without any prior knowledge. However, most of the learned latent variables remain unexplained by the available descriptors. Yet, the model’s performance implies that these learned representations are *useful* in affinity prediction. This suggests that there are both similarities and differences in how machines “see” chemicals versus how human experts see them. Understanding this distinction may further improve model performance or reveal new mechanisms behind drug– target interactions.

Meanwhile, the distribution of the test set errors suggest that there are “problem drugs” (and “problem proteins”) for which prediction is especially difficult. One could action this insight either by collecting more training data for these drugs (or proteins), or by using domain-knowledge to engineer features that complement the molecular graphs. Indeed, knowing that the PCA outliers are the easiest to predict suggests that some additional feature input may be needed to differentiate between drugs that lack distinct molecular sub-graphs. Although 2D graphs contain more information than 1D strings, our model still neglects the stereochemistry of the molecules. Future experiments could test whether representing drugs in 3D (or proteins in 2D) further improves model performance.

Interestingly, under-representation of proteins in the training set does seem to be the reason for the “problem proteins”. The **Supplement**^3^ shows an analysis of the effect of homologous proteins on test set performance. Although we see that test set error varies across clustered protein groups, the training set represents all protein clusters equally well. This suggests that the variation in test set performance is not simply explained by asymmetrical representation of protein groups within the training set.

## 4 Summary and Future Work

We test **GraphDTA** with four graph neural network variants, including GCN, GAT, GIN, and a combined GAT-GCN architecture, for the task of drug–affinity prediction. We benchmark the performance of these models on the Davis and Kiba datasets. We find **GraphDTA** performs well for two separate benchmark datasets and for two key performance metrics. In a post-hoc statistical analysis of our model, we find that **GraphDTA** can learn the importance of known molecular descriptors without any prior knowledge. We also examine the model’s performance and find that a handful of drugs contribute disproportionately to the total prediction error. Although we focus on drug–target affinity prediction, our **GraphDTA** model is a generic solution for any similar problem where either data input can be represented as a graph.

It may be possible to improve performance further by representing proteins as graphs too, for example by a graph of their 3D structure. However, determining the 3D structure of a target protein is very challenging. We chose to use the primary protein sequence because it is readily available. The use of 1D sequences, instead of 3D structures, also reduces the number of parameters that we need to learn, making it less likely that we over-fit our model to the training data. Still, for some problem applications, it may make sense to use structural information, as well as binding site, binding confirmation, and solution environment information, to augment the model.

## Footnotes

1 https://www.synapse.org/#!Synapse:syn15667962/wiki/583305

2 https://www.synapse.org/#!Synapse:syn15667962/wiki/592145

3 https://github.com/thinng/GraphDTA/blob/master/supplement.pdf

